# An integrated single-cell transcriptome landscape of postnatal mouse hypothalamus

**DOI:** 10.1101/2023.04.13.536811

**Authors:** Muhammad Junaid, Han Kyoung Choe, Kunio Kondoh, Eun Jeong Lee, Su Bin Lim

**Author notes:** **To whom correspondence should be addressed.** Su Bin Lim, Department of Biochemistry & Molecular Biology, Ajou University School of Medicine, Suwon 16499, Korea., Eun Jeong Lee, Department of Brain Science, Ajou University School of Medicine, Suwon 16499, Korea.

## Abstract

The neural stem cells (NSCs) in the hypothalamus are relatively more narrowly defined than in other neurogenic regions of the postnatal brain. By leveraging single-cell RNA sequencing (scRNA-seq) data, we generated an integrated reference dataset comprising 296,282 cells from postnatal hypothalamic regions and the adjacent region, bed nucleus of the stria terminalis (BNST), and identified 30 hypothalamic neuronal and non-neuronal cell populations. The analyses of their gene expression pattern, and specific differentiation trajectories reveal the presence of NSCs and intermediate progenitor cells (IPCs) that show a continuum of activation and differentiation processes in the hypothalamus after birth. Through comparative analyses of the integrated dataset with our lab-generated Connect-seq data obtained from the whole hypothalamus, we further assessed the technical validity of the dataset presented in this study. Our large-scale unified scRNA-seq dataset with harmonized cell-level metadata can serve as a valuable resource for investigating cell type-specific gene expression and cellular differentiation trajectories in the postnatal mouse hypothalamus.

## Introduction

Postnatal neurogenesis involves precursor cell division, differentiation, and the integration of newly formed cells into the brain^1,2^. Over the past decade, studies have provided increasing evidence that neural stem cells (NSCs) may be responsible for the bulk of postnatal neurogenesis^3,4^. These stem cells can produce both neuronal progenitor cells (NPCs) and glial progenitor cells, which in turn differentiate into postmitotic neurons and glial cells, respectively.

In recent years, studies have demonstrated that neurogenesis/gliogenesis can also occur in the adult mammalian hypothalamus^5,6^. Although the hypothalamus is a critical brain region that plays a vital role in regulating homeostatic and survival-related behaviors, there is still limited knowledge about its intrinsic mechanisms of development^7-9^. Despite many years of study, the existence and extent of adult neurogenesis/gliogenesis remain a topic of debate. One possible reason for such discrepancy could be a lack of well-characterized sub-populations of cells in the hypothalamus that are specifically pivotal for this process. Studies that have used techniques such as immunochemistry and in situ hybridization have been limited by the number of transcripts and markers available, thereby challenging the identification and assessment of these cell populations and their locations^10^. A comprehensive analysis of the populations of NSCs and progenitor cells throughout the entire hypothalamus is thus required to gain a deeper understanding of the cell proliferation in the hypothalamus of postnatal animals.

The high-throughput single-cell RNA sequencing (scRNA-seq) analytical and experimental approaches have been applied to study the proliferation of the brain in depth^11,12^. Particularly, the hypothalamic region has been extensively studied in regard to neurogenesis/gliogenesis and development^13^. A variety of proliferating cell types has been identified in this region, including glial-like cells, neuronal progenitors, astrocyte progenitors, and oligodendrocyte progenitors^14-16^. More recently, scRNA-seq data analysis of 207,785 cells of adult macaque hippocampus revealed a comprehensive single-cell transcriptome atlas of neurogenesis for primates^10^. However, cellular and molecular characterization of postnatal mouse hypothalamic neurogenesis/gliogenesis with single-cell resolution is still lacking, possibly due to limited large-scale scRNA-seq dataset with uniform cell-level metadata containing major and minor cell types and other biologically relevant information^17^.

Our goal is to identify and extensively characterize the cell-type-specific features during neurogenesis/gliogenesis in the hypothalamic region of mice in an unprecedented resolution, from postnatal stages of development. We processed and analyzed publicly available scRNA-seq transcriptomic data that were obtained from the hypothalamic regions of mice, using uniform and optimized informatics pipeline. Here, we performed an integrated single-cell transcriptome analysis of the postnatal mouse hypothalamus, comprising 294,749 cells, and analysed cell differentiation trajectories and properties during neurogenesis/gliogenesis (Fig. 1). Our integrated dataset has revealed 30 distinct cell types that encompass all major cell types found in the hypothalamic regions, including glial-like cells such as ependymal cells, tanycytes, astrocytes, oligodendrocytes, and intermediate progenitor cells (IPCs). Additionally, we found co-expression of the neural precursor marker *Sox2* and the proliferating marker *Mki67*, which confirms the presence of NSCs in the postnatal and young adult mouse hypothalamus^10^. Our data analysis has revealed the transcriptional dynamics that underlie oligodendrocytes differentiation and cellular heterogeneity of tanycytes, a type of non-neuronal cell that is specific to the hypothalamus and whose function in the adult brain is not well understood. Furthermore, to assess the technical validity of our integrated dataset, we performed anchor-based mapping and transferring of labels using our lab-generated Connect-seq-derived single nuclei RNA-seq (snRNA-seq) dataset of 1,533 cells from whole hypothalamus and the bed nucleus of the stria terminalis (BNST) of CRH-IRES-Cre (CRH-Cre) mice. The presented large-scale single-cell transcriptomic data can serve as a valuable asset for exploring gene expression and cellular differentiation in the hypothalamus across various stages of development.

**Figure 1.**
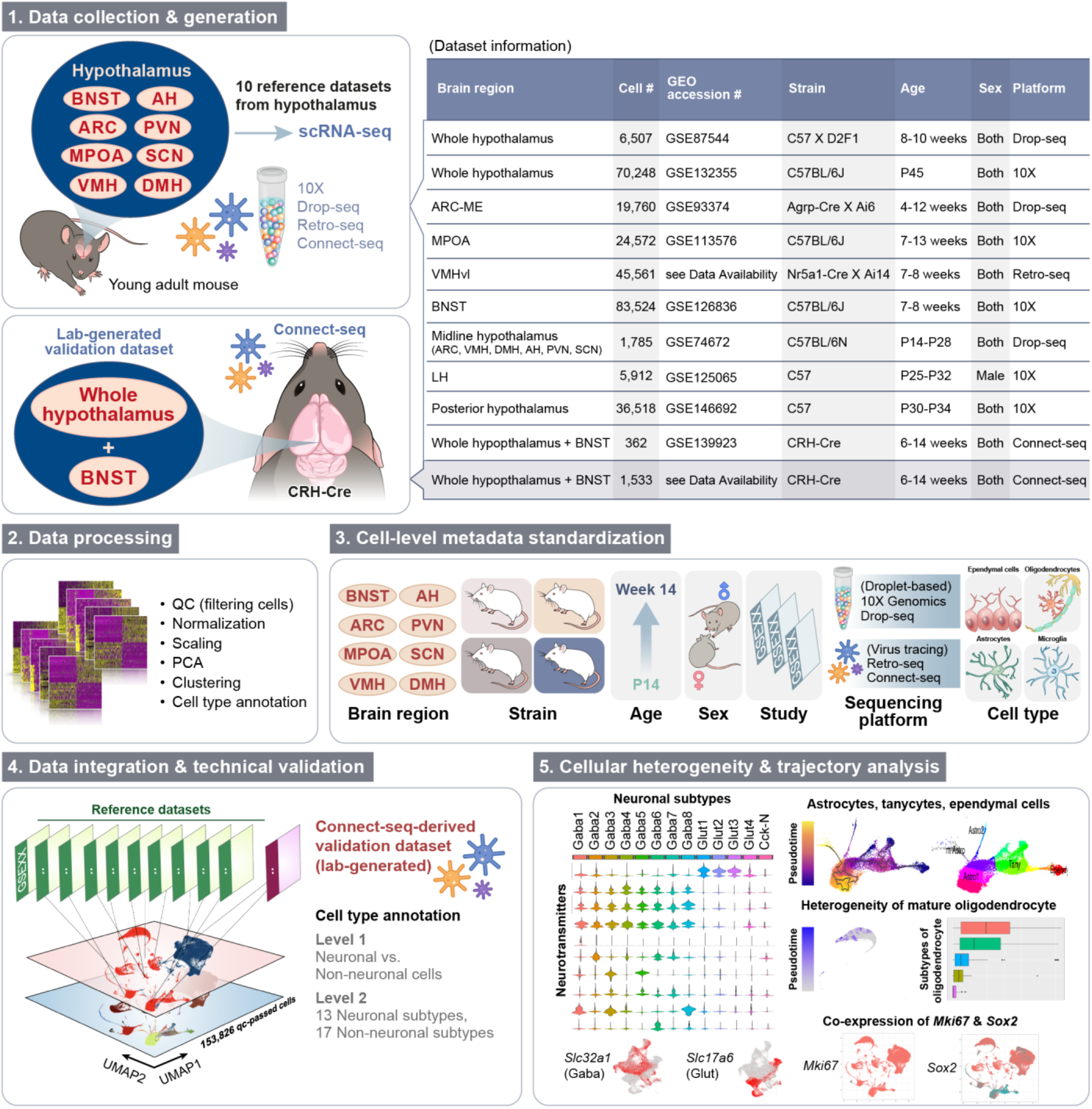
Schematic representation of the study design, batch correction, and clustering for single-cell RNA-seq of young adult mouse brain datasets.

## Results

### Generation and analysis of integrated scRNA-seq data of postnatal hypothalamic regions

We collected ten publicly available scRNA-seq datasets of mice from various hypothalamic brain regions including arcuate hypothalamus and median eminence (ARC-ME), lateral hypothalamus (LH), medial preoptic area/anterior hypothalamus (MPOA), midline hypothalamus (Mid-hypo), ventral posterior hypothalamus (VPH), ventromedial hypothalamus (VMHvl), and the bed nucleus of the stria terminalis (BNST), with ages ranging from P14 to 14 weeks, from the NCBI GEO. (see Methods and Data Availability). The datasets were obtained from mice of various strains including C57BL/6, C57 X D2F1, Agrp-Cre X Ai6, Nr5a1-Cre X Ai14, and CRH-Cre^8,14,18-24^. Here, we used a uniform informatics pipeline for re-analysis, including quality control, standardization of cell-level metadata, mapping, and label transferring against the validation dataset (Fig. 1). To facilitate re-use of the integrated data, we manually curated cell-level attributes, such as study, age, brain region, sex, and strain of mouse, of each dataset and used standardized texts for metadata. After integrating and correcting for technical batch effects that might arise from different studies and experimental protocols, we performed principal component (PC) analysis (Supplementary Fig. 1) and dimensionality reduction using uniform manifold approximation and projection (UMAP). Mitochondria-related read counts and unique feature counts were used to filter a total of 294,749 cells of the integrated data (Fig. 2A), in which 152,524 QC-passed cells were subsequently clustered into 18 cell populations using cell-type specific markers in the first iteration of clustering. Cells were distributed relatively evenly throughout the dataset and major cell type (Fig. 2B). To determine if the virus tracing-based sequencing approaches, such as Retro-seq^25,26^ and Connect-seq^24^, would identify cells with varying proportions compared to more commonly used droplet-based tools such as 10X^27^ and Drop-seq^28^, we computed cell proportion by major cell types across different sequencing platforms (Fig. 2C). While Retro-seq-derived dataset had relatively fewer QC-passed neurons and more oligodendrocytes and astrocytes, the relative proportions of each cell type of Connect-seq-derived dataset were comparable between cells of datasets obtained from droplet-based sequencing platforms. Fig. 2D shows the QC-passed cells colored by different cell-level attributes before and after batch removal.

**Figure 2.**
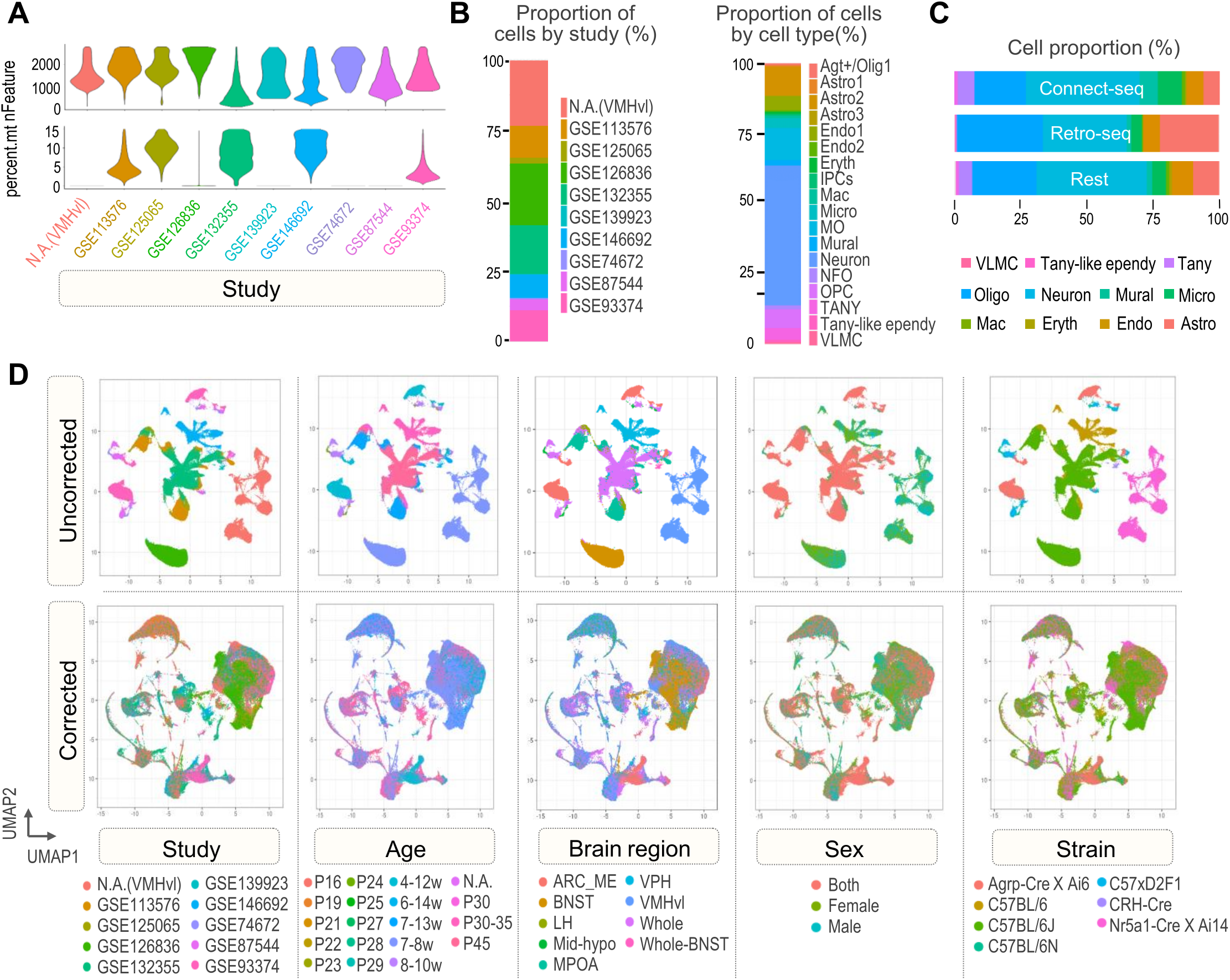
Cell type proportion and batch removal of final merged dataset. (A) Cells of the final merged dataset are grouped by cell types, including percentage of read counts from mitochondrial genes (percent.mt), number of unique genes (nFeature). (B) Proportions of cell number are grouped by study (left) and cell type (right), which are further divided by sequencing tool. (D) UMAP plots of the batch uncorrected (top) and corrected (bottom) merged dataset. Cells are grouped by study, age, brain region, sex, and mouse strain.

We first distinguished neuronal cells from non-neuronal cells using the known pan-neural genes (*Snap25, Syp, Elval2, Slc17a6*, and *Slc32a1*) and canonical non-neural genes (*Olig1, Cldn5, C1qa, Sox9*, and *Mustn1*)^14,23,29,30^ (Fig. 3A-B). To characterize diverse populations of neurons in the postnatal hypothalamic regions, we next examined the expression patterns of markers for excitatory and inhibitory neurons, which are *Slc17a6* for glutamatergic neurons and *Slc32a1*, Gad1 and Gad2 for GABAergic neurons, respectively (Fig. 3C)^31^. Of the 13 identified neuronal clusters, there are 4 glutamatergic clusters, 8 GABAergic clusters and one cluster neither glutamatergic nor GABAergic, but *Cck*+ (Fig. C-D). Overall, within the identified neuronal cell populations, clusters can be identified by the unique combination of genes that are expressed at different levels (Fig. 3E).

**Figure 3.**
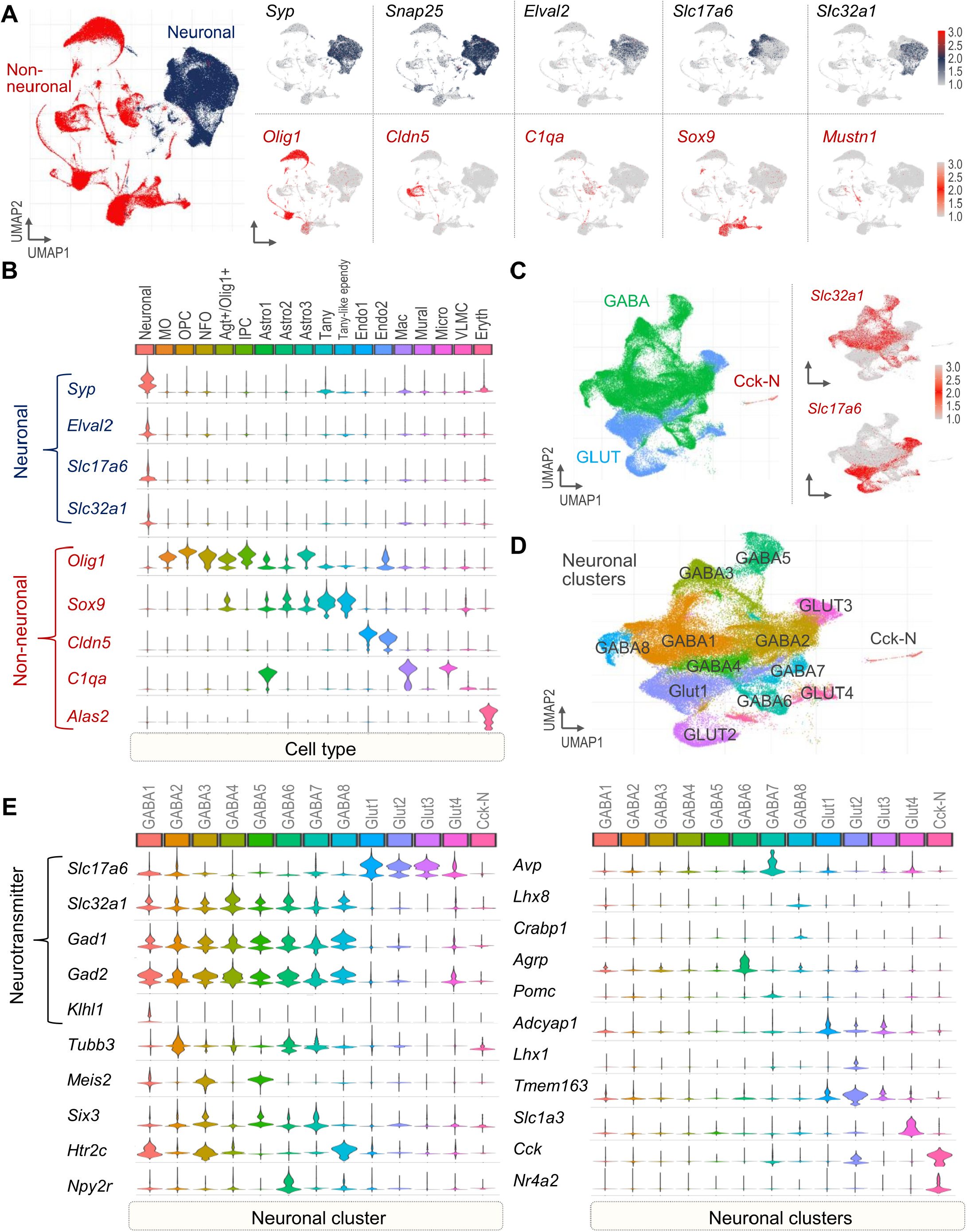
Cell type annotation and analyses of the merged dataset of young adult mouse hypothalamus. (A) UMAP plots showing neuronal or non-neuronal cells (left) classified by combined expression of of pan-neuronal markers (right). (B) Relative expression of pan-neuronal and non-neuronal marker genes are shown. (C) UMAP plots showing normalized expression of *Slc17a6, Slc32a1* after the second iteration of unsupervised clustering on just neuronal cells. (D) UMAP plot showing 13 neuronal cell types identified by unsupervised clustering. (E) Relative expression of neurotransmitters (*Slc17a6, Slc32a1, Gad1, Gad2*) and discriminatory marker genes.

Next, we identified a total of 17 distinct clusters of non-neural cells (Fig. 4A-B). Oligodendrocytes were found using markers, such as *Olig1, Pdgfra, Fyn*, and *Mobp*, all of which show a gradient of gene expression in postnatal development^14,23,32,33^ (Fig. 4A-B and Supplementary Figs. 2 and 3). These cells also include oligodendrocytes progenitor cells (OPC), newly formed oligodendrocytes (NFO), and myelinating oligodendrocytes (MO). We also identified three distinct subtypes of astrocytes (Astro1, 2, and 3) expressing *Agt* at varying levels. The Astro1 cell cluster exhibits high expression of *Gfap*, while the cells of Astro2 highly express of *Unc13c* and *Rgcc*. The Astro3 cell cluster has high expression of *C1qa* and low expression of *Unc13c*^34,35^ (Fig. 4C). Tanycytes (Tany) and tanycyte-like ependymal cell (Tany-like ependy) were also found using the expression levels of *Rax* and *Ccdc153*, respectively. In addition, we found other distinct clusters readily identifiable as macrophages (Mac: *Mrc1*^*+*^), microglia (Micro: *Cx3cr1*^*+*^), two populations of endothelial cells (Endo1/2: *Pecam1*^*+*^), vascular leptomeningeal cells or VLMCs (*Dcn*^*+*^) and one erythrocyte representing cluster (Eryth: *Alas2*^*+*^) (Fig. 4A-B). These findings of major non-neural cells obtained from the hypothalamus at varying postnatal stages closely align with previous scRNA-seq analyses from mouse brains^23,24,36,37^.

**Figure 4.**
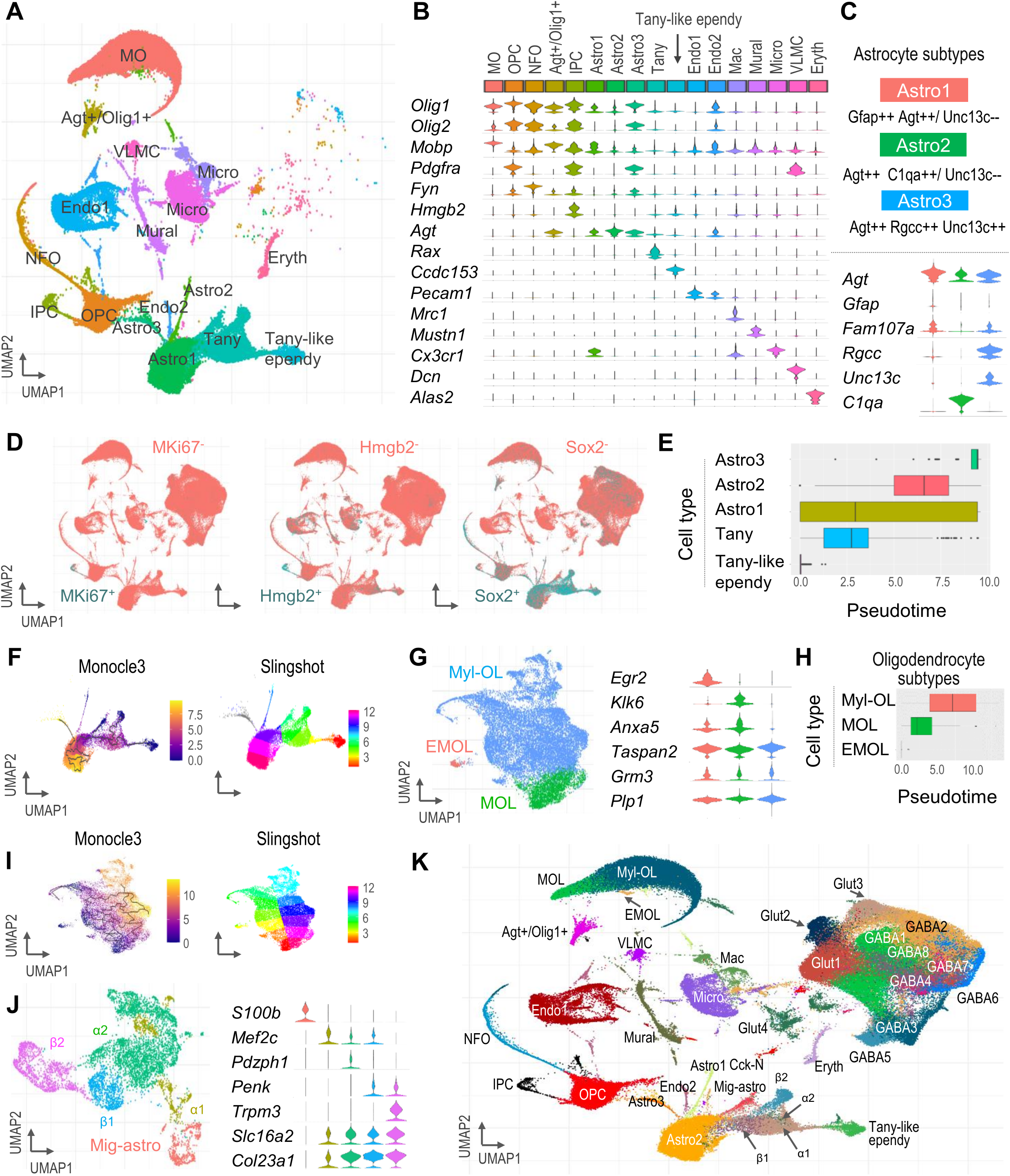
Cell type annotation and analyses of non-neuronal cell clusters. (A) UMAP plot showing 17 non-neuronal cell types identified by unsupervised clustering. (B) Relative expression of cell type-specific genes differentially expressed across the 17 non-neuronal cell subtypes. (C) Relative expression of astrocyte subtype-specific genes. Positive and negative markers for each subtype are indicated by “++” and “--”, respectively. (D) UMAP plots showing positive and negative expression of Mki67/Hmgb2/Sox2 genes across all cell types in young adult hypothalamus. (E,F) Pseudotime ordering of cells determined by Monocle3 and slingshot and slinghot showing a single trajectory from “Tany-like ependy” to “Astro3” cells. (G) UMAP plot (left) and relative expression (right) of oligodendrocyte subtype-specific genes showing the heterogeneity of MO clusters after the second iteration of unsupervised clustering. (H, I) Pseudotime ordering of cells determined by Monocle3 and slingshot and slinghot showing a single trajectory and heterogeneity in MO differentiation. (J) UMAP plot (left) and relative expression (right) of tanycyte subtype-specific genes in combined hypothalamic tanycytes. (K) UMAP plot showing subtypes of neuronal and non-neuronal cells.

### Subpopulations of glial cells

We next aimed to identify and characterize subpopulations of glial cells in the hypothalamus. Tany-like ependymal cells, tanycytes, and astrocytes express both canonical marker gene *Gfap*, the neural precursor marker gene *Sox2* as well as *Sox9*, which encodes the transcription factor required for the maintenance of glial progenitor cells^38^ (Fig. 4D, Supplementary Fig. 2A-B). To study the process of differentiation and specialization of these glial-like cell populations, we applied subtyping (Supplementary Fig. 2A) and trajectory analysis tools using the monocle3 and slingshot (see Methods), and found that pseudotime ordering of cells increased from ‘Tany-like ependy’ cells to mature astrocytes (Astro3), suggesting differentiation of tanycyte-like ependymal cells into astrocytes in the postnatal/ adult hypothalamus (Fig. 4E-F and Supplementary Fig. 2A). Interestingly, we further detected a subpopulation of *Olig2*^+^ astrocytes (Supplementary Fig. 2C), supporting a recent immunohistochemistry study that has discovered Olig2-expressing astrocytes that are highly enriched in specific regions in adult mouse brain, in which their function in postnatal or adult hypothalamushas yet to be found^39^.

### Subpopulations of proliferating progenitor cells

We also found two proliferating progenitors’ populations (IPCs and OPCs). IPCs are a population of NSCs and play a crucial role in adult neurogenesis in the hypothalamus by differentiating into excitatory and/or inhibitory neurons^40^. IPCs can be identified by the expression of *Mki67* (a marker gene for proliferation), *Sox2* (a marker gene for neural precursor cell), and *Hmgb2*, which encodes a chromatin-related transcription activator that is crucial for the transition of NSCs from quiescence to proliferative status^10,41^ (Fig. 4B and D). These subpopulations of cells expressing *Sox9, Sox2*, and *Mki67/Hmgb2* in the hypothalamic regions may undergo the cell division process, before, during, and after the nuclear division, as suggested by recent studies that have found the presence of such proliferative NSCs in postnatal hypothalamic regions^42,43^. OPCs are another type of neural progenitor cell that give rise to oligodendrocytes^44,45^. Notably, these cells express *Pdgfra* (Fig. 4B), which regulates processes of cell proliferation, differentiation and migration through the Wnt and Notch signalling pathways, indicating their potential role in regulation of adult oligogenesis in postnatal hypothalamus.

### Subpopulations of oligodendrocytes

Subtyping of myelinating oligodendrocytes (MO) revealed 3 clusters: (1) early mature oligodendrocytes (EMOL) expressing *Egr2* (Early Growth Response 2; also known as *Krox20*) during oligodendrocytes maturation, (2) mature oligodendrocytes (MOL) expressing late oligodendrocyte differentiation genes, such as *Klk6 and Anxa5*, and (3) myelin-forming oligodendrocytes (Myl-OL) expressing genes involved in myelin formation, such as Ctps, *Mog, Plp1, and Mobp*^46,47^ (Fig. 4G and Supplementary Fig. 3B-C). Trajectory analyses of oligodendrocyte populations and MO subtypes revealed intermediate stages of differentiation (Fig. 4H-I and Supplementary Fig.3A-C), in line with the previously reported subpopulations of oligodendrocytes expressing intermediate expression of *Olig2* and/or *Sox2*^48^.

### Subpopulations of tanycytes

In the hypothalamus, tanycytes and associated cells of the ventricular zone are strong candidate stem/progenitor cells that express transcription factors such as *Rax, Sox2, Sox9*, and genes encoding intermediate filament proteins including GFAP (Supplementary Fig. 2B). To study transcriptional heterogeneity of tanycyte subtypes, we divided tanycytes into four major subtypes – *α1, α2, β1*, and *β2* – based on their gene expression using subtype-specific markers, as shown in Fig. 4J. Interestingly, we found one cell cluster expressing *S100b*, which is known to be involved in the regulation and migration of astrocytes^49^. Altogether, our integrated large-scale scRNA-seq dataset with standardized cell-level metadata including annotations of major and minor cell populations (Fig. 4K) may help to advance the understanding of transcriptome dynamics of neuronal and non-neuronal cells including proliferative precursor cells with differentiation potencies in the hypothalamus of postnatal and young adult mouse.

### Generation of validation dataset

To further assess technical validity of our integrated data (‘reference’ dataset), we generated and analyzed Connect-seq-derived single nuclei RNA-seq (snRNA-seq) dataset comprising 1,533 cells (‘validation’ dataset) with previously applied processing workflow (see Methods, Fig. 5A and Supplementary Fig. 4A-C). Using known marker genes, we identified a total of 8 cell clusters, including IPCs (*Mki67*^+^), OPC (*Pdgfra*^*+*^), Astro (*Agt*^*+*^), Endo1 (*Pecam*^*+*^), Endo2 (*Itm2a*^*+*^), Mac (*C1qa*^*+*^), MO (*Mog*^*+*^) and one neuronal cluster expressing *Syp* (Fig. 5B-C). By applying an anchor-based multi-modal reference mapping (see Methods), we transferred the cell type classifications of the integrated dataset onto the validation dataset (Fig. 5A), leading to generation of ‘predicted’ annotations (Fig. 5D). The integrated-mapped dataset helps us to identify cell types that were previously blended in an [unsupervised analysis of the validation dataset](https://satijalab.org/seurat/articles/pbmc3k_tutorial.html) (Supplementary Fig. 5). The integrated and validation datasets were merged into a final dataset comprising 153,826 cells with standardized cell-level attributes including study, sex, mouse strain, age, brain region, and cell type (Fig. 5E-F).

**Figure 5.**
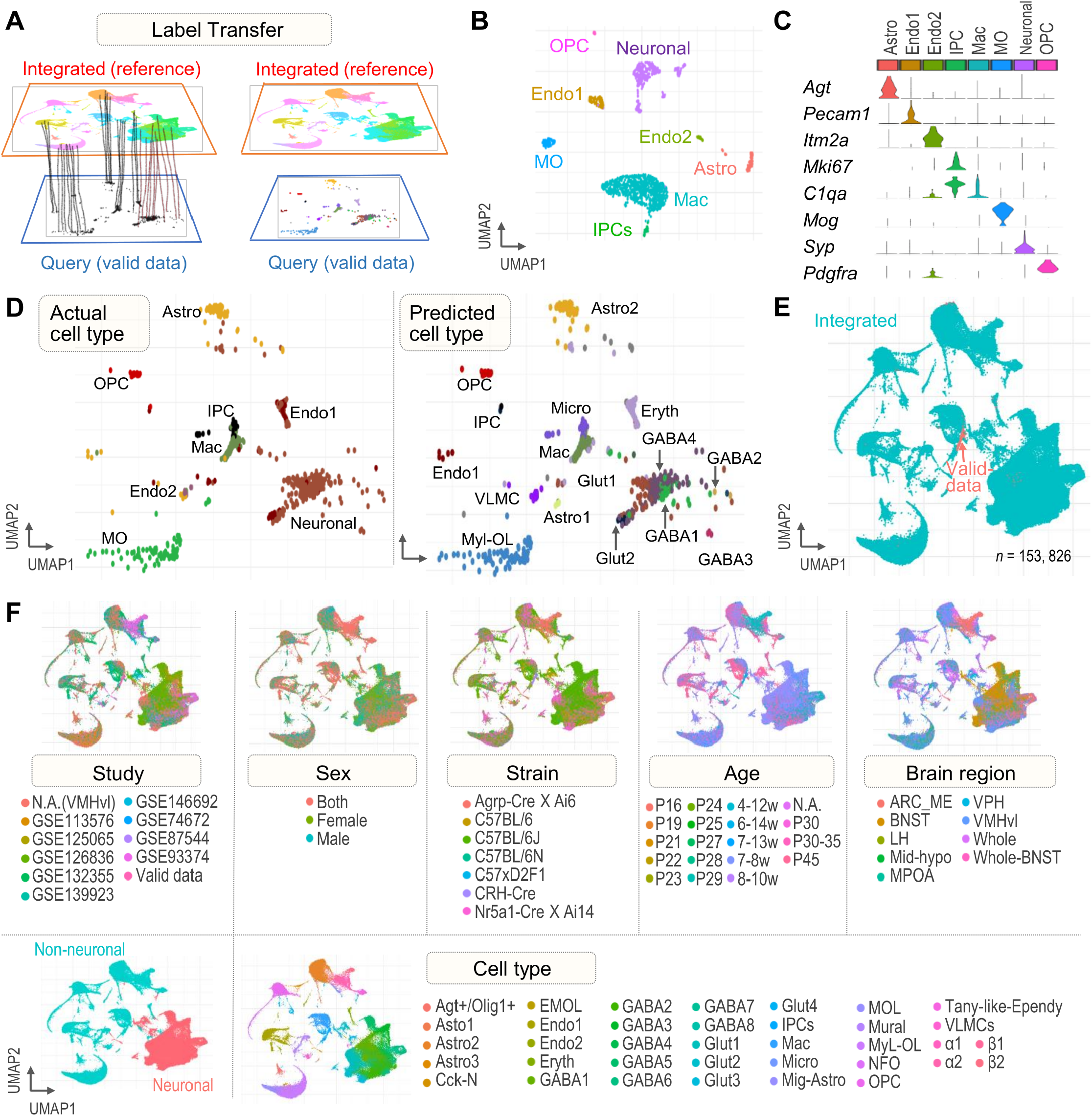
Mapping and transferring labels of merged dataset for technical validation. (A) Schematic representation showing identified anchors, which facilitate transfer of discrete labels between integrated (reference) and query (lab-generated validation) dataset. (B) UMAP plot showing the cell type annotation of validation dataset (C) Relative expression of neuronal and non-neuronal gene markers. (D) UMAP plots showing the actual and predicted cell types on validation dataset by projecting cells of validation dataset onto UMAP embeddings of the integrated dataset. (E) UMAP plot showing cells of the validation dataset and the integrated dataset in the final UMAP embeddings. (F) UMAP plots of the final merged (reference and validation) dataset. Cells are grouped by study, sex, mouse strain, age, brain region, and cell type.

## Discussion

*In vitro* studies have demonstrated that NSCs can both self-renew and differentiate into various cell types, validating functional tri-potency of NSCs (i.e., the ability to generate neurons, oligodendrocytes, and astrocytes)^50-52^. Our presented large-scale scRNA-seq dataset may further advance the understanding of diverse cell types and their transcriptional dynamics underlying NSC lineages, differentiation and specialization processes. We identified key subpopulations of NSCs and NPCs that may be indicative of neurogenesis and gliogenesis occurring in postnatal and young adult mouse brain^10,53,54^. The existence of actively growing glial cells and IPCs suggests the sturdy occurrence of neurogenesis during the postnatal period of development, particularly in the adult hippocampus^55,56^. In the hypothalamus, glial cells have been identified as potential NSCs that play important and specific roles in regulating various physiological processes such as osmolarity, circadian rhythm, metabolism, and cell proliferation. In order to gain insights into the molecular mechanisms that control the differentiation and specialization of glial cells in the hypothalamus during postnatal and adult neurogenesis/gliogenesis, we have identified and characterized each population of glial-like cells. By utilizing the high-throughput integrated dataset, we were able to validate the progenitor cell populations from two perspectives: (1) the expression of canonical marker genes and (2) the differentiation direction inferred by the pseudo-time and slingshot analysis.

Very recently, Steuernagel et al. presented a reference map of the murine hypothalamus consisting of 384,925 cells by integrating public scRNA-seq datasets, including those obtained from various stages including the embryonic stage^57^. This unified atlas successfully defines a variety of physiologically relevant neuronal cell subtypes with harmonized annotation, but postnatal to adult hypothalamic neurogenesis/gliogenesis still requires more in-depth molecular characterization. It is not yet known how adult animals maintain their NSC pools. The existence of a population of self-renewing glial cells that prevents NSC depletion is a topic of ongoing debate^58-61^. In this study, we specifically identified key NSC and NPCs cell types in postnatal hypothalamic regions. We found that the expression of *Hmgb2, Mki67*, and *Sox9/Sox2* genes that are associated with proliferation may play a role in this process. Furthermore, the expression of the *Mki67* and *Hmgb2* genes in IPCs suggests that these specific subpopulations of progenitor cells may actively undergo proliferation. The presence of the Sox2/Sox9^+^ cells, which are markers of glial cells further indicate the existence of proliferative NSCs within this population. In addition, we applied an anchor-based mapping method to Connect-seq-derived snRNA-seq dataset with our integrated reference dataset to predict cell type annotations and further confirm the similarity of transcriptional profiles between the two datasets. The advances in experimental and analytical tools for single-cell sequencing research will continue to uncover previously unknown cell subpopulations with functional roles and reveal lineage dynamics of NSCs in postnatal and adult mouse brain, providing a more comprehensive understanding of adult stem cell lineages^62-65^.

## Acknowledgements

We acknowledge support provided by the Korea Health Technology R&D Project through the Korea Health Industry Development Institute (KHIDI), funded by the Ministry of Health & Welfare, Republic of Korea (HR22C1734) and the National Research Foundation (NRF) of Korea (2020R1A6A1A03043539, 2020M3A9D8037604, 2022R1C1C1004756). E.J.L is supported by the National Research Foundation (NRF) of Korea (2022R1C1C1005741) and the new faculty research fund of Ajou University School of Medicine.

## Author contributions

S.B.L. and E.J.L. conceptualized and designed the study. E.J.L. and H.K.C generated the validation dataset. K.K provided PRV. M.J., S.B.L., and E.J.L. analyzed and interpreted the data. All authors reviewed and contributed to the manuscript.

## STAR*METHODS

### KEY RESOURCES TABLE

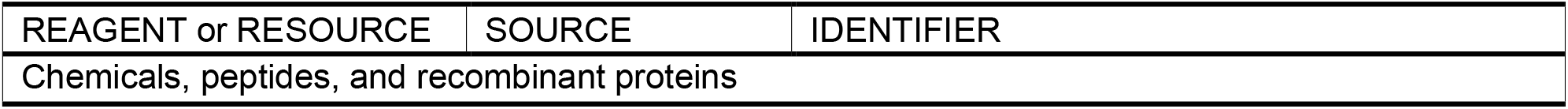
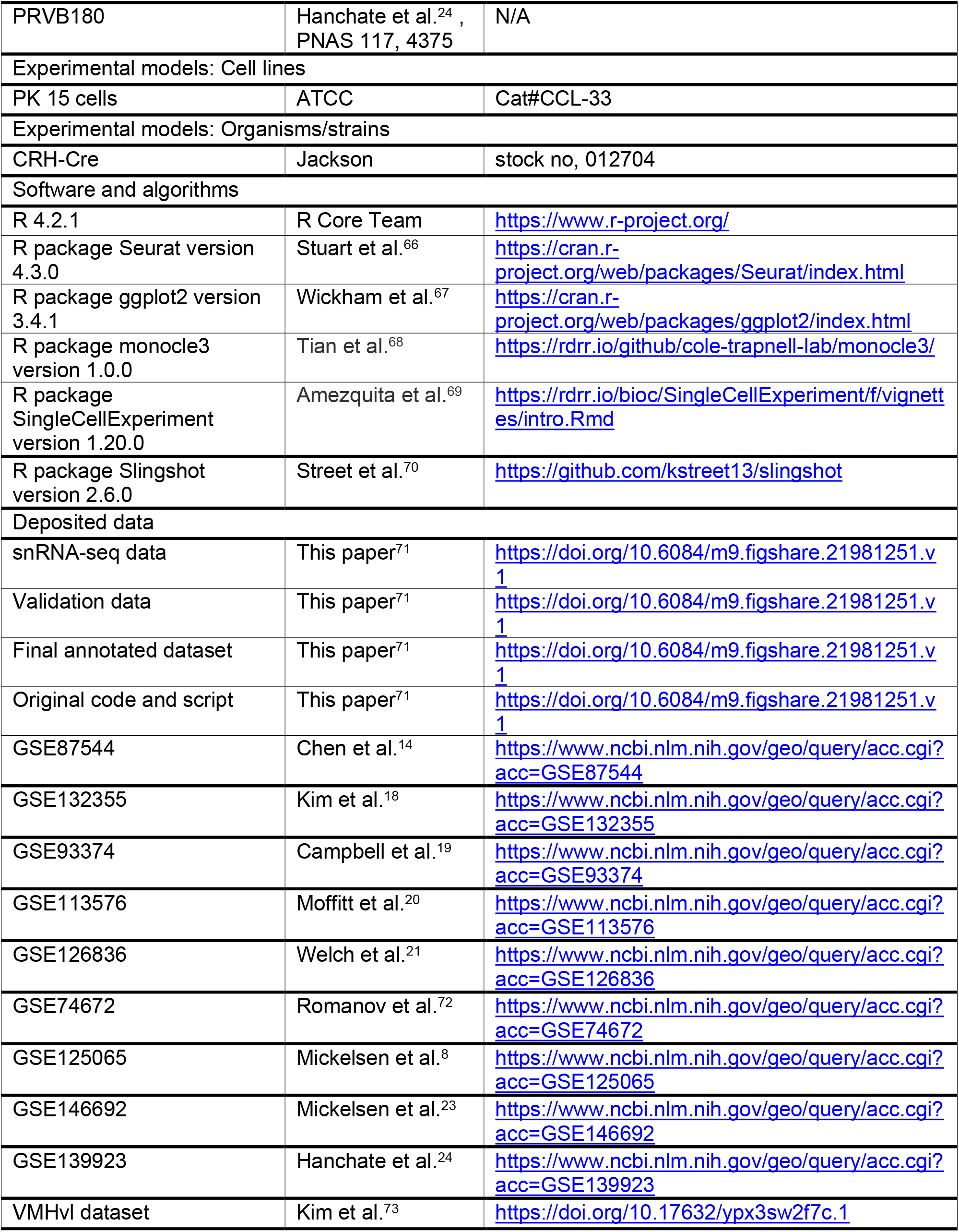

## Resource availability

### Lead contact

Further information and requests for resources and reagents should be directed to and will be fulfilled by the lead contact, Prof. Su Bin Lim and Prof. Eun Jeong Lee.

### Materials availability

This study did not generate new unique reagents.

## Experimental model and subject details

### Mice

To produce validation data, CRH-IRES-Cre (CRH-Cre; stock no, 012704) mice were purchased form The Jackson Laboratory. All procedures involving mice were approved by the Ajou University Institutional Animal Care and Use Committee and DGIST Institutional Animal Care and Use Committee.

## Method details

### PRV (pseudorabies virus)

PRVB180 was propagated following methods described previously^24,74^. Briefly, to propagate PRV180, PK15 cells (American Type Culture Collection) were infected with the virus using a multiplicity of infection of 0.1 to 0.01. After infection, cells showed a prominent cytopathic effect (∼2 days). They were harvested by scraping, and the cell material was frozen using liquid nitrogen and then quickly thawed in a 37°C water bath. After three freeze-thaw cycles, cell debris was removed by centrifugation twice at 1000g for 5 min, and the supernatant was then used for experiments. The titer of viral stocks was determined using standard plaque assays on PK15 cells^75^, with titers expressed in plaque-forming units (pfu).

### Stereotaxic Injections

Viruses were injected into the PVN of CRH-Cre mice as described previously^24^. All injections were done under inhalation anesthesia of 2% isoflurane. Briefly, 1 μL of PRVs (PRVB180; 1 to 1.5 × 106 p.f.u.) were loaded into a 1-μL syringe and injected bilaterally into the brain at a rate of 100 nL/min using a Stereotaxic Alignment System (David Kopf Instruments). The needle was inserted into the PVN based on a stereotaxic atlas^76^. After recovery, animals were singly housed with regular 12-h dark/light cycles, and food and water were provided ad libitum.

### Processing and integration of scRNA-seq datasets

We obtained 10 scRNA-seq datasets from independent studies using from the NCBI GEO, which were further processed using the Seurat package^77^ in R (v. 4.2.1). A standard informatics pipeline was applied to each dataset for pre-processing and cell clustering for cell-level metadata standardization. Cells that have more than 15% of mitochondria-related read counts and unique feature counts over 2,500 or less than 100 were filtered out. A total of 152,524 cells was included for further analysis, and variable features of these cells were identified after normalization. The FindIntegrateionAnchors() function of the Seurat was applied to merged cells to find common anchors among variables. The number of principal components (PCs) of the dataset was determined using the Elbow plot prior to clustering of cells (Supplementary Fig. 1). Cells of the merged dataset were clustered into distinct cell populations using the FindClusters() function (resolution = 0.25). The resulting Louvain clusters were visualized in a two-dimensional UMAP representation and were manually annotated with known cell types using canonical neural and non-neural marker genes, as well as signatures chosen in the original publications.

### Characterization and analysis of cell subpopulations

To identify the various populations of excitatory and inhibitory neurons, we subsetted the neuronal cell cluster of the merged data and performed second round of dimensional reduction and clustering using the FindVariableFeatures(), ScaleData(), RunPCA(), and RunUMAP() functions of the Seurat package (resolution = 0.3). Distinct clusters of neuronal cells were annotated with known marker genes, such as *Slc17a6, Slc32a1, Cck, Gad1 and Gad2*. Similarly, tanycytes and mature oligodendrocytes were subsetted to run second round of dimensional reduction and clustering and thus to characterize cellular heterogeneity and pseudotime trajectory.

### Pseudotime analysis

Pseudotime trajectory analyses of ‘Tany-like ependy’, ‘Tany’, ‘Astro1’, ‘Astro2’, ‘Astro3’, ‘Myl-OL’, ‘MOL’, and ‘EMOL’ cells were performed using the Monocle3^78-80^ Alpha package in R (v. 1.0.0). A standard Monocle workflow was applied to these cells using applying the importCDS() and reduceDimension() functions, which perform normalization, dimensionality reduction, visualization, and differential expression analyses by computing the size factors that control for variability in library construction efficiency among cells. In addition, the slingshot package in R (v. 2.6.0)^70,81,82^ was used to validate Monocle3-derived pseudotime trajectories. To get UMAP trajectory interference based on the pseudotime values the “slingshot” function was used that shows the relative position of each cell along the trajectory.

### Isolation of single nuclei

Connect-seq experiments were performed as previously described^24^ with modifications for nuclei isolations, Adult 8-week-old CRH-Cre mice were injected with PRVB180. After 3 days, mice were euthanized by cervical dislocation, and the brain quickly removed. The hypothalamus was carefully microdissected under a microscope and immediately placed into a nuclei isolation medium (sucrose 0.25 M, KCl 25 mM, MgCl2 5 mM, TrisCl 10 mM, dithiothreitol, 0.1 % Triton). Tissue was Dounce homogenized, allowing for mechanical separation of nuclei from cells^83^. The nucleic acid stain Hoechst 33342 (5 μM, Life Technologies) was included in the media to facilitate visualization of the nuclei. Samples were washed, resuspended in nuclei storage buffer (sucrose, MgCl2 5 mM, and TrisCl 10 mM) and filtered. Solutions and samples were kept cold throughout the protocol. Then, PRV-infected single nuclei were isolated based on the fluorescence emitted by TK-GFP using flow cytometry (FACSAria II; BD Biosciences) according to methods described previously^24^.

### Single nuclei capture

FACS-sorted single nuclei were loaded onto an independent single channel of a Chromium Controller (10× Genomics) single-cell platform. Briefly, 10,000 single nuclei were loaded for capture using a Chromium Single Cell 3′ Reagent kit, v2 Chemistry (10× Genomics). Following capture and lysis, complementary DNA was synthesized and amplified (14 cycles) as per the manufacturer’s protocol (10× Genomics). The amplified cDNA was used to construct an Illumina sequencing library and sequenced on a single lane of a NovaSeq 6000 (Illumina). For FASTQ generation and alignments, Illumina basecall files (*.bcl) were converted to FASTQs using Cell Ranger v.1.3 (10× Genomics), which uses bcl2fastq v.2.17.1.14. FASTQ files were then aligned to mm10 genome and transcriptome using the Cell Ranger v.1.3 pipeline, which generates a ‘gene vs cell’ expression matrix.

### Connect-seq-derived data

This dataset was used for validation purpose to assess the technical validity of the integrated dataset, and was processed using the uniform informatics pipeline, as previously. For cell clustering, resolution was set as 0.25, and 8 cell clusters were identified using specific marker genes, such as *Agt, Pecam1, Itm2a, Mki67, C1qa, Mog, Syp*, and *Pdgfra*. These manually annotated cell types were compared with the cell types predicted by the ‘referenced-based’ integration and ‘label transfer’ method from the Seurat, using the FindTransferAnchors() and TransferData() functions, respectively (Fig. 5D and Supplementary Fig. 5 Fig. 5). Next, we filtered out the cells of validation dataset having predicted cell type score (‘predicted.celltype.score’) less than 0.5, and used the remaining cells to merge with the integrated dataset and to visualize their cell-level attributes in final UMAP (Fig. 5E, F).

## Data and code availability

The R script used to generate the integrated dataset, conduct validation dataset analysis, and produce relevant plots is available in Figshare (DOI:https://doi.org/10.6084/m9.figshare.21981251.v1)^71^. Single-cell RNA-seq datasets used to generate the ‘integrated’ datasets are available at the NCBI GEO under the accession numbers GSE87544^14^, GSE132355^18^, GSE93374^19^, GSE113576^20^, GSE126836^21^, GSE74672^72,84,85^, GSE125065^8^, GSE146692^23^, GSE139923^24^ and a scRNA-seq dataset of VMHvl hypothalamic region can be accessed at https://doi.org/10.17632/ypx3sw2f7c.1 (Mendeley Data)^73^. Our final dataset containing both integration and validation datasets can be found in Figshare (https://doi.org/10.6084/m9.figshare.21981251.v1)^71^.

## Supplemental information

Figure S1. QC and processing of the integrated data. (A, B) Heatmap and VizDimReduction plots showing the distribution and ordering of both cells and features according to their PCA scores. (C) Elbowplot shows the ranking of principle components based on the percentage of variance explained by each one.

Figure S2. Sox9 expressing glial-like cells (A) UMAP plot shows proposed neurogenic differentiation pathway from previously published literature (B) Feature plots shows some neurogenic gene expression across Sox9 expressing cells (C) Violin plot shoes Olig2-expressing astrocyte subtype heterogeneity (Olig2-AS).

Figure S3. Transcriptional heterogeneity expression during oligodendrocyte Maturation. (A) Box plots showing pseudotime ordering of IPCs (blue), OPCs (green), NFOs (Cyan), and MOs (red) based on their gene expression profiles. (B) Pseudotime gene expression analysis shows the gene-specific heterogeneity expression of the MO cluster. (C) Expression patterns of stage-specific genes during oligodendrocyte maturation.

Figure S4. QC and processing of the validataion dataset. (A, B) Heatmap and VizDimReduction plots showing the distribution and ordering of both cells and features according to their PCA scores. (C) Elbowplot shows the ranking of principle components based on the percentage of variance explained by each one.

Figure S5. Feature plots showing predicted cell type on validation dataset after projecting cells onto integrated UMAPs.

