## Supplementary Figures for "An integrated single-cell transcriptome landscape of postnatal mouse hypothalamus"

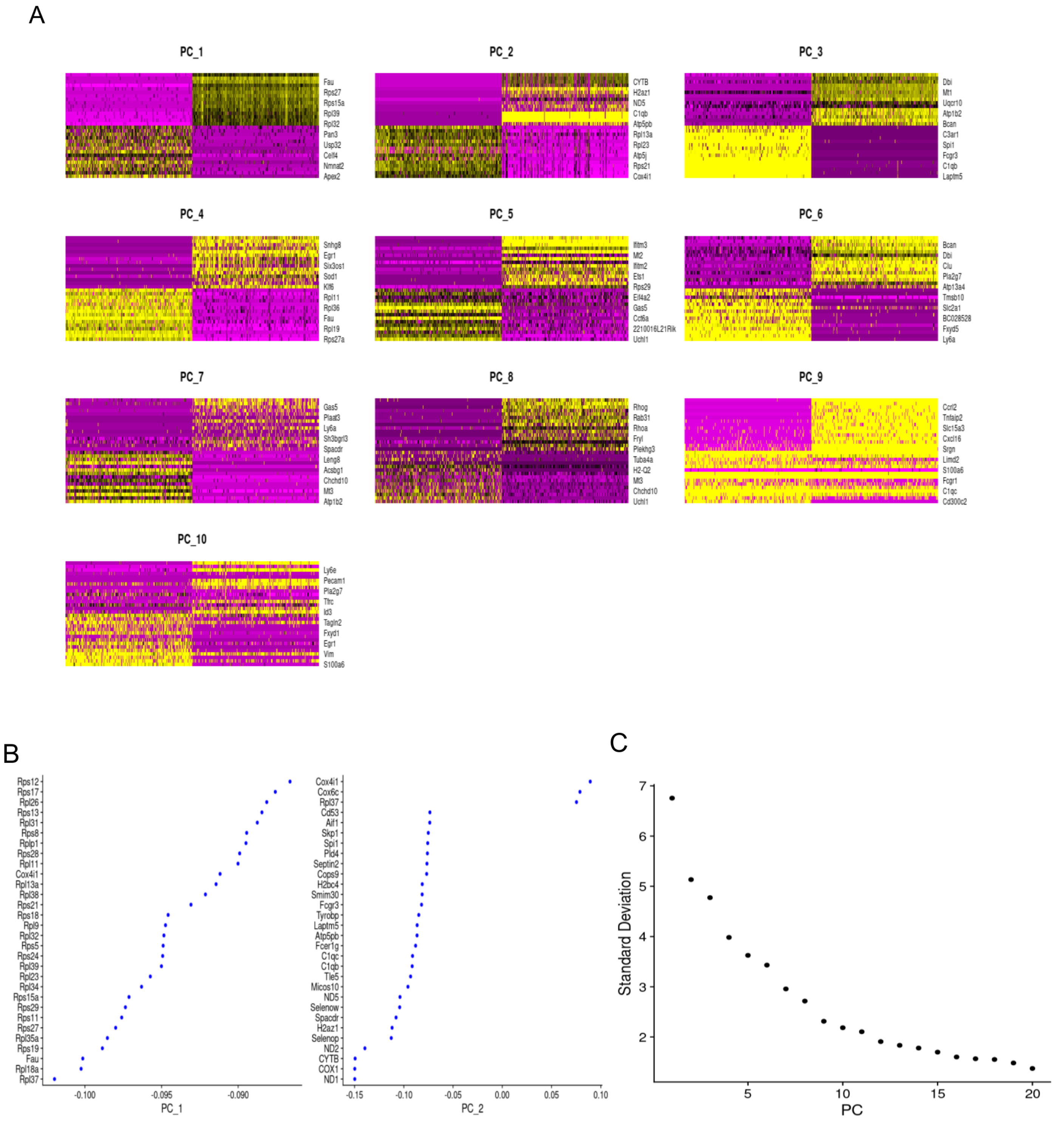


**Figure S1.** QC and processing of the integrated data. (A, B) Heatmap and VizDimReduction plots showing the distribution and ordering of both cells and features according to their PCA scores. (C) Elbowplot shows the ranking of principle components based on the percentage of variance explained by each one.

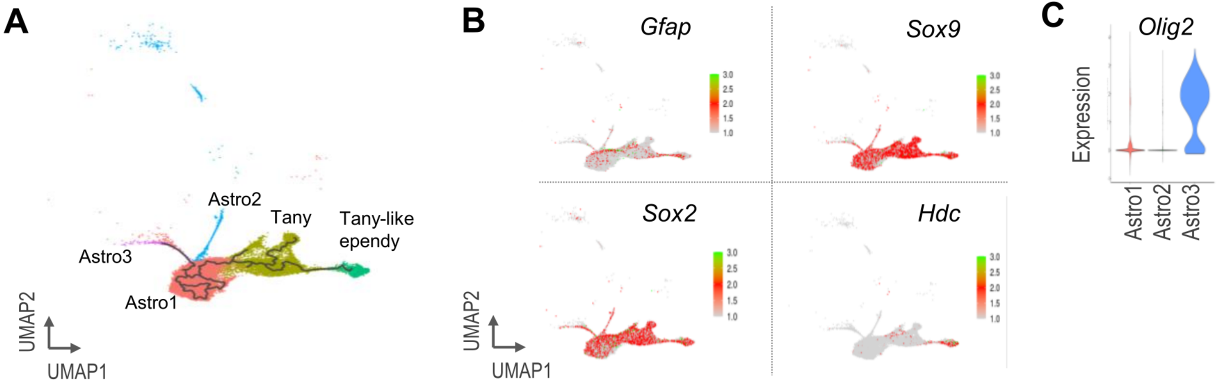


**Figure S2.** Neurogenesis related gene expression analysis Sox9 expressing glial-like cells (A) UMAP plot shows proposed neurogenic differentiation pathway from previously published literature (B) Feature plots shows some neurogenic gene expression across Sox9 expressing cells (C) Violin plot shoes Olig2-expressing astrocyte subtype heterogeneity (Olig2-AS).

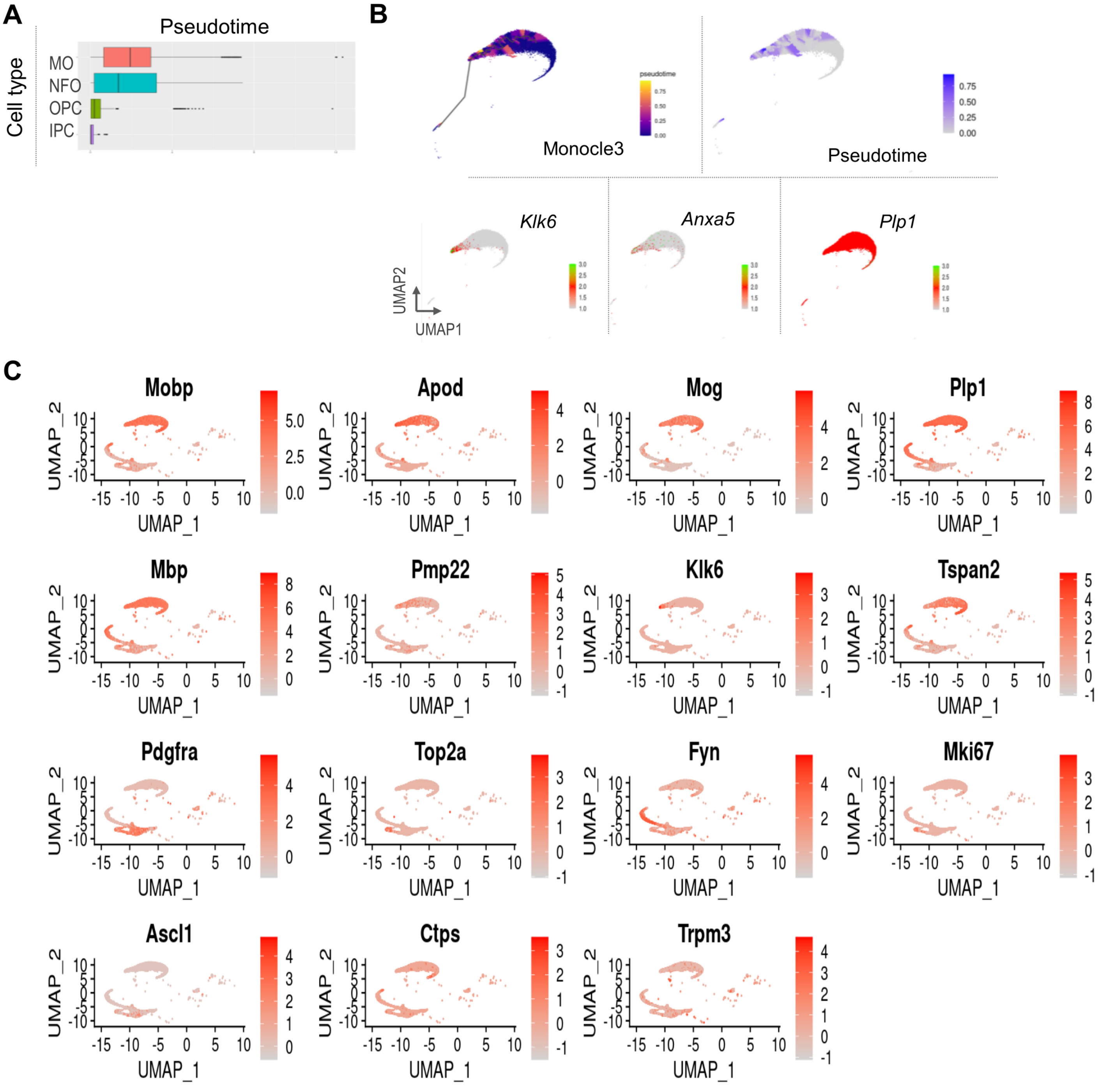


**Figure S3.** Transcriptional heterogeneity expression during Oligodendrocyte Maturation. (A) Box plots showing pseudotime ordering of ordering of IPCs (blue), OPCs (green), NFOs (Cyan), and MOs (red) based on their gene expression profiles. (B) Pseudotime gene expression analysis shows the gene specific heterogeneity expression of MO cluster. (C) Expression patterns of stage-specific genes during oligodendrocyte maturation

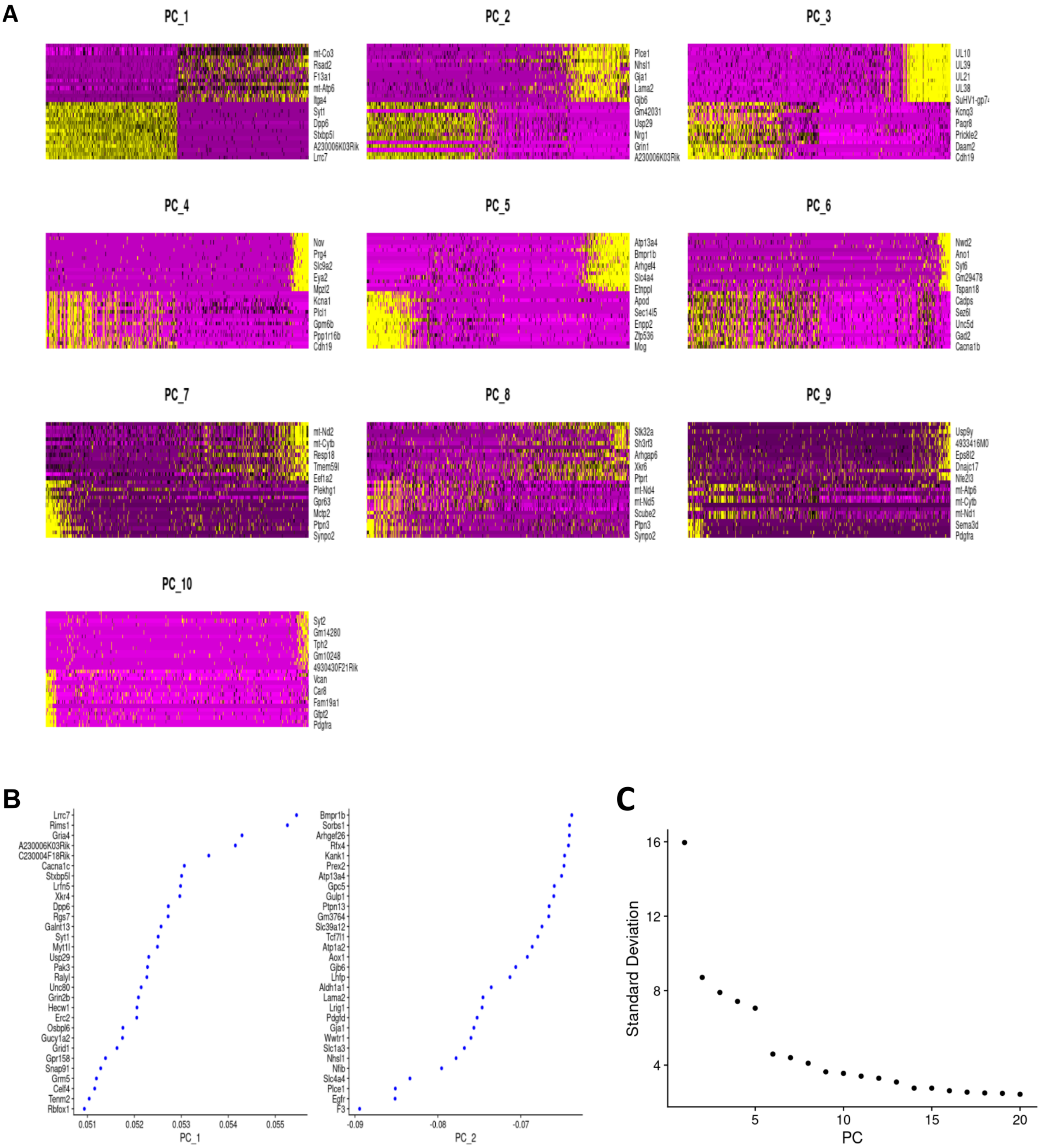


**Figure S4.** QC and processing of the validation dataset. (A, B) Heatmap and VizDimReduction plots show the distribution and ordering of both cells and features according to their PCA scores. (C) Elbowplot shows the ranking of principal components based on the percentage of variance explained by each one.

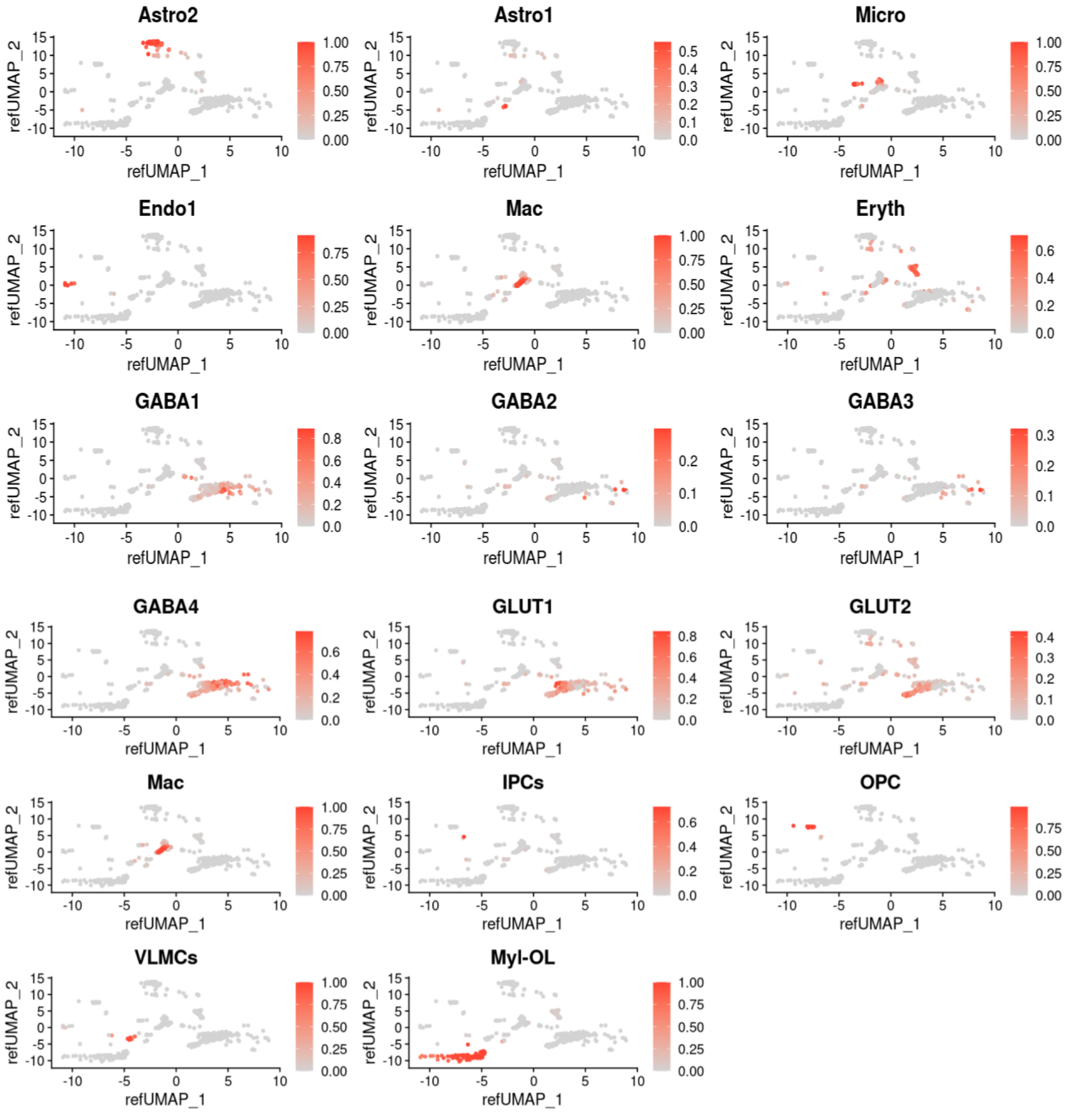


**Figure S5.** Feature plots showing predicted cell type on validation dataset after projecting cells onto integrated UMAPs.
